# TNAP limits TGF-β-dependent cardiac and skeletal muscle fibrosis by inactivating SMAD2/3 transcription factors

**DOI:** 10.1101/655332

**Authors:** Benedetta Arnò, Francesco Galli, Urmas Roostalu, Bashar Aldeiri, Tetsuaki Miyake, Alessandra Albertini, Laricia Bragg, Sukhpal Prehar, John C. McDermott, Elizabeth J. Cartwright, Giulio Cossu

**Author notes:** Correspondence to: Prof. Giulio Cossu Division of Cell Matrix Biology & Regenerative Medicine Faculty of Biology, Medicine and Health, University of Manchester. Manchester Academic Health Science Centre Michael Smith Building, D.4316 Oxford Road. M13 9PL Manchester, UK. These authors contributed equally to the work (Benedetta Arnò, Francesco Galli).

## Abstract

Fibrosis is associated with almost all forms of chronic cardiac and skeletal muscle diseases. The accumulation of extracellular matrix impairs the contractility of muscle cells contributing to organ failure. Transforming growth factor beta (TGF-β) plays a pivotal role in fibrosis, activating pro-fibrotic gene programs via phosphorylation of SMAD2/3 transcription factors. However, the mechanisms that control de-phosphorylation of SMAD2/3 have remained poorly characterized. Here we show that tissue non-specific alkaline phosphatase (TNAP) is highly upregulated in hypertrophic hearts and in dystrophic skeletal muscles, and the abrogation of TGF-β signalling in TNAP positive cells reduces vascular and interstitial fibrosis. We show that TNAP co-localizes and interacts with SMAD2. TNAP inhibitor MLS-0038949 increases SMAD2/3 phosphorylation, while TNAP overexpression reduces SMAD2/3 phosphorylation and the expression of downstream fibrotic genes. Overall our data demonstrate that TNAP negatively regulates TGF-β signalling and likely represents a mechanism to limit fibrosis.

**Summary statement:** This paper shows that tissue non-specific alkaline phosphatase negatively regulates TGF-β signalling and may represent a mechanism to limit fibrosis through SMAD dephosphorylation.

## Introduction

Fibrosis accompanies many chronic diseases and is a common response to injury. It is characterized by excessive production and accumulation of collagen and other extracellular matrix (ECM) components, cellular dysfunction and loss of tissue architecture that eventually lead to organ failure (Ueha et al., 2012). Fibrosis can affect many organs, including heart and skeletal muscle, and has become a major cause of death in the developed world.

Fibrosis is an integral component of most cardiac pathological conditions and can be associated with cardiomyocyte death (Frangogiannis, 2012), pressure or volume overload (Berk et al., 2007), hypertrophic cardiomyopathy (Kania et al., 2009) and toxic insults (Bernaba et al., 2010).

Skeletal muscle fibrosis accompanies aging, in which case gradual muscle loss leads to the accumulation of adipose tissue and ECM (Alnaqeeb et al., 1984; Wood et al., 2014). Fibrosis occurs early in congenital muscular dystrophies and impairs muscle function and regeneration (Serrano and Munoz-Canoves, 2017; Tedesco and Cossu, 2012). In both cardiac and skeletal muscle, fibrosis increases the stiffness of the tissue, thereby directly impairing muscle cell contraction.

The molecular pathways involved in fibrosis are well known and common for the two organs. Transforming Growth Factor beta (TGF-β), in addition to being a major morphogen during development, is one of the main signaling molecules initiating fibrosis and is secreted by many cell types in the injured tissue. Its activity is controlled by proteolytic cleavage of a precursor, eventually bound to inhibitory proteins and subsequently released by proteases. Active TGF-β binds to the serine threonine protein kinases receptors (termed TGF-β receptor type I and type II) and induces the phosphorylation and activation of SMAD2/3 transcription factors (also known as receptor regulated SMADs or R-SMADs) (Heldin et al., 1997; Zi et al., 2012). The phosphorylation of R-SMADs triggers their association with SMAD4, leading to their nuclear translocation, where they control the transcription of pro-fibrotic genes. Finally, the dephosphorylation of R-SMADs promotes their exit from the nucleus and the end of the signaling (Bruce and Sapkota, 2012). As a central regulator of a number of physiologically important processes TGFβ signaling is highly regulated. Cellular susceptibility is determined by receptor endocytosis via clathrin and caveolin dependent pathways (Di Guglielmo et al., 2003). Depletion of TGFβ ligand defines the transcriptional response and is achieved by TGFβ binding to the cell surface or its cellular uptake (Clarke et al., 2009). The duration and temporal dynamics of ligand availability influence downstream signaling response, as repeated pulses of TGFβ can lead to long-term sustained response (Zi et al., 2011). Several post-translational modifications are capable of regulating R-SMAD activity. These include ubiquitinylation (Dupont et al., 2012; Dupont et al., 2005; Episkopou et al., 2001; Lin et al., 2000; Zhu et al., 1999), acetylation (Gronroos et al., 2002), parpylation (Lonn et al., 2010), sumoylation (Miles et al., 2008) and phosphorylation at different serine residues (Kretzschmar et al., 1997). Rapid inactivation of R-SMADs can be achieved by dephosphorylation. However, to date only one R-SMAD phosphatase, PPM1A (Protein Phosphatase 1A), has been identified for the TGF-β pathway (Lin et al., 2006). Identification of new R-SMAD phosphatases would enable efficient attenuation of pro-fibrotic signaling in acute and chronic diseases.

Tissue Non Specific Alkaline Phosphatase (TNAP) is an enzyme widely expressed in many organs. It is involved in bone mineralization and mutations in the TNAP gene have been associated with hypophosphatasia, a rare inherited disorder, characterized by defective bone mineralization (Mornet et al., 2001). Recently TNAP has been associated with vascular calcification (Romanelli et al., 2017; Sheen et al., 2015) and cardiac fibro-calcification (Herencia et al., 2015). Its role in fibrosis and in the context of TGF-β signaling has not been addressed.

In the present study we demonstrate that TNAP negatively regulates TGF-β signaling pathway by regulating the level of phosphorylation of SMAD 2/3. We show that TNAP is highly induced in hypertrophic hearts and dystrophic skeletal muscle and the abrogation of TGF-β signaling in TNAP positive cells reduces vascular and interstitial fibrosis. TNAP interacts with SMAD2 and its overexpression decreases SMAD phosphorylation and the expression of downstream fibrotic genes.

## Results

We previously established TNAP expression in skeletal muscle and demonstrated that its high activity marks perivascular cells (Dellavalle et al., 2007). In contrast, we found that TNAP is more widely expressed in healthy mouse heart. We detected alkaline phosphatase activity mainly in the interstitium (Figure 1a,b) but also in cardiomyocytes (Figure 1c). At higher resolution (Figure 1d) AP activity co-localized both with endothelial (VE-Cadherin +) and perivascular (NG2+) cells and also extended outside of the small vessels (fig. 1d). Of the three AP isoforms, Intestinal, Embryonic and Non Specific, only the latter is expressed in the heart at any developmental stage examined (Fig. 1e). When TNAP-Cre^ERT^ mice (Dellavalle et al. 2011) were crossed to NGZ reporter mice, LacZ activity was detected in many areas of the heart (Figure 1f). Additional examples of TNAP localization in adult healthy heart are shown in Supplementary Figure 1.

**Figure 1.**
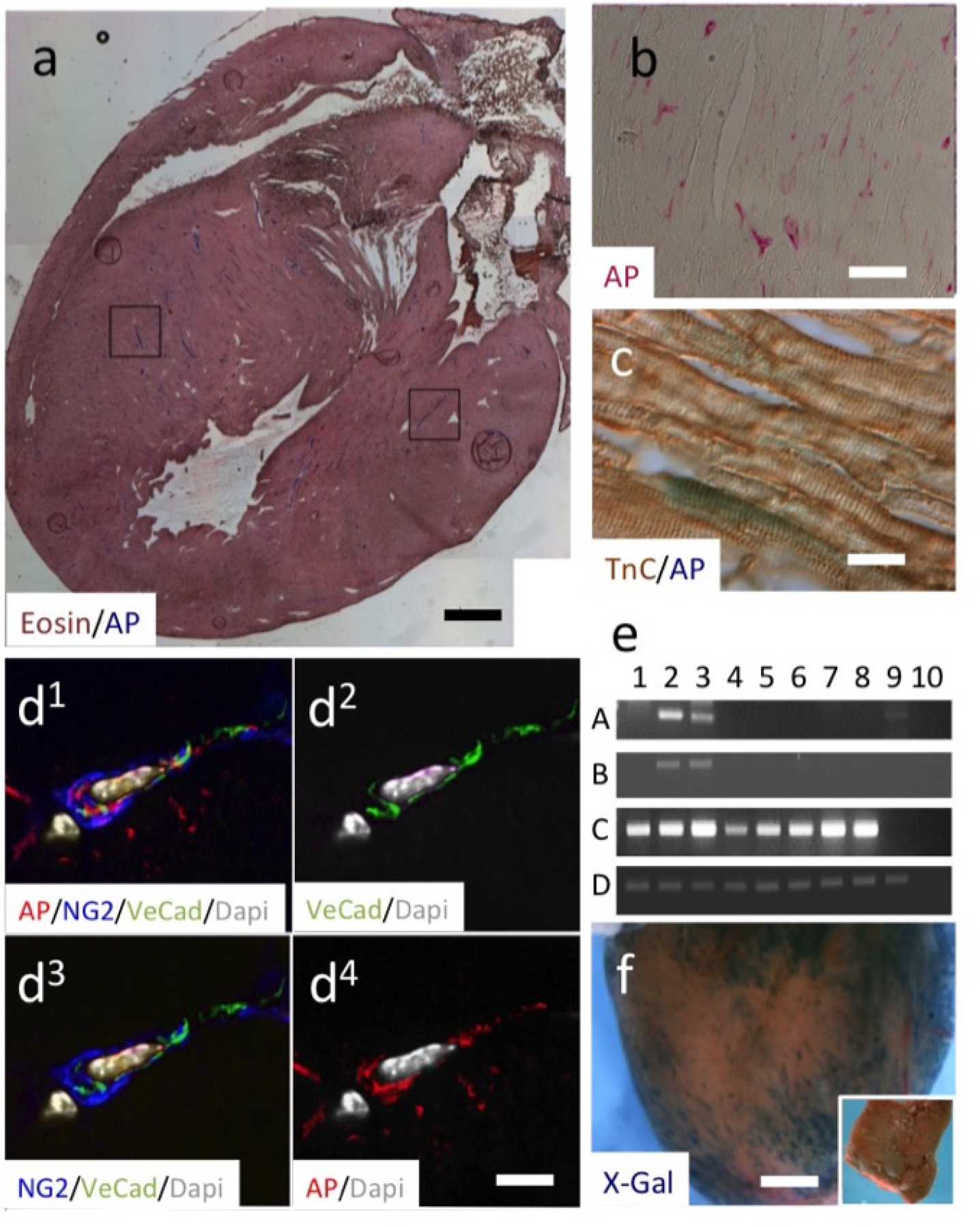
TNAP activity under healthy, physiological conditions. (a) Frontal section of an adult (P30) mouse heart stained for alkaline phosphatase activity and eosin. Alkaline phosphatase activity is high small and medium size vessel (squares in Fig. 1a and Fig. 2b) and is also observed in some cardiomyocyte (Fig. 1c) (d) Confocal triple immunofluorescence of a small vessel stained with anti-VE Cadherin (green) and anti-NG2 (blue) antibodies and for AP activity (red). (e) RT-PCR with probes specific for embryonic (A), intestinal (B), tissue non specific (C) alkaline phosphatase and for GAPDH (D). 1: adult liver; 2: adult intestine; 3: adult testis; 4:embryonic heart (E 11.5); 5: foetal heart (E 16.5); 6: neonatal heart (P1); 7: juvenile heart (P15); 8: adult heart (P90); 9 D16 (embryonic mesoangioblasts: Minasi et al. 2002); 10: negative control. (f) TNAP activity is maintained in adulthood. Tamoxifen was injected for 3 consecutive days in TNAP-Cre^ERT^ x Rosa NGZ at P6,7,8 and the heart was collected at P60 (Dellavalle et al. 2011). A control injection of oil is shown in the inset. Scale bars: a 1mm, b,c, 50 μm; d, 15 μm; f, 0.5 mm.

### Angiotensin II infusion increases TNAP expression in cardiomyocytes and cardiac fibroblast in vivo

We next analysed changes in TNAP activity in response to fibrosis. We used a model of Angiotensin II-induced cardiac hypertrophy, which is known to cause fibrosis after few days (Crowley et al., 2006). We implanted subcutaneous mini-pumps releasing Angiotensin II (Ang II) (2.8mg/kg per day) for two weeks in 10 weeks old C57/BL6J mice. We analysed the hearts of treated mice for the presence of hypertrophy and fibrosis. As expected, the treatment with Ang II caused cardiomyocytes hypertrophy and led to an increase in Tenascin C positive areas compared with mice treated with saline solution (Sham) (Supplementary Fig. 2a-i). Moreover, the expression of the fibrotic genes *Fibronectin1, Tenascin C, Sma2, Vimentin* and *Collagen1a2* was significantly upregulated in Ang II treated mice compared with the control (Supplementary Fig. 2j-p).

**Figure 2.**
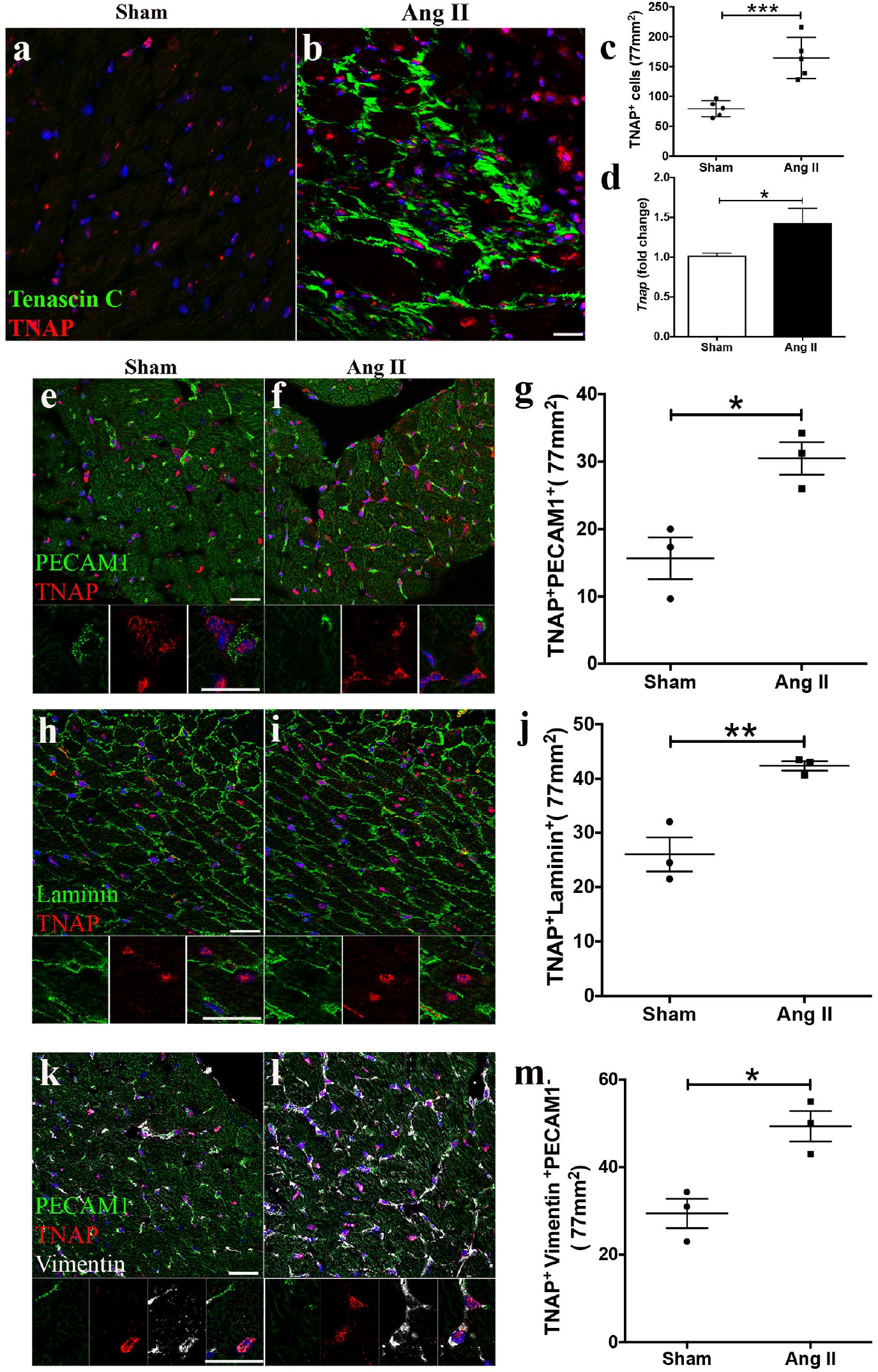
TNAP is upregulated in different cell types in Ang II-induced fibrosis. (a,b) Immunofluorescence for Tenascin C and TNAP in Sham and Ang II treated mice (c) Quantification of TNAP positive cells on both samples (mean ± S.D., n= 5). (d) RT-qPCR for Tnap in Sham and Ang II treated ventricle extracts (mean ± S.D., n=5). Confocal images of Sham and Ang II treated heart sections stained for TNAP and PECAM1 (e, f); TNAP and Laminin (h, i); TNAP, PECAM1 and Vimentin (k, l). Insets show high magnifications of double positive cells. Quantifications of double positive cells are indicated in g, j and m respectively (mean ± S.D., n=3). Scale bars: 20 µm. A two-tailed t-test was used to assess statistical significance: * p<0.05, ** p<0.01, *** p<0.001.

We took advantage of this disease model to analyse the expression of *Tnap*. Interestingly, we found a significant increase in the number of TNAP^+^ cells in hypertrophic hearts in comparison to the control (Fig. 2a-c; n=5 for each group). Moreover, we detected an upregulation of *Tnap* expression by RT-qPCR (Figure 2d). In order to understand which cell types express TNAP in physiological and pathological conditions, we stained the heart sections for TNAP and Laminin, Vimentin and PECAM, which allow identification of cardiomyocytes basal lamina, cardiac interstitial and endothelial cells respectively, and using confocal microscopy counted the number of double positive cells in a 77 mm^2^ area. We confirmed that all three cells types analysed expressed TNAP in normal heart (TNAP/PECAM^+^ cells 16.6 ± 6%, TNAP/Laminin^+^ cells 26 ± 5.4%, TNAP/Vimentin^+^/PECAM^−^ cells 27 ± 5.6%, n=3 mice, mean ± s.d.). Remarkably, in the Ang II treated mice the number of cells expressing TNAP significantly increased in all three different cell populations (TNAP^+^/PECAM^+^ cells 30.5 ± 4.1%; TNAP^+^/Laminin^+^ cells 42.4 ± 1.5%, TNAP^+^/Vimentin^+^/PECAM^-^ cell;s 49.3 ± 6%, n=3 mice, mean ± s.d., P<0.001, two-tailed t-test) (Fig. 2e-m). These data prove that in hypertrophic hearts TNAP is significantly up regulated in cardiac myocytes, endothelial cells and interstitial cells. However, it is important to notice that, under normal condition only a minority of cells in the heart express TNAP and even if this number almost doubles in Ang treated animal, it remains confined to less than half of total cardiac cells.

### The abrogation of TGF-β signaling in TNAP positive cells reduces fibrosis in the heart and skeletal muscle

Since the increased number of TNAP^+^ cells is preferentially localised in Tenascin C positive regions (Fig. 2a,b,k,l), we reasoned that TNAP^+^ cells might have a role in the regulation of fibrosis upon Ang II treatment. TNAP is widely expressed in many different cells so that a cardiac specific knock out would be technically very challenging in this context as Cre should be driven by al least three (cardiac, endothelial and interstitial) different promoters. Therefore, to prove that TNAP expressing cells are in fact involved in cardiac fibrosis, despite being a minority of total cardiac cells, we inactivated TGF-β signalling specifically in TNAP positive cells. To achieve this goal we took advantage of an inducible conditional knockout *Tnap*^*cre*^;*Tgfβr2* floxed mouse (Chytil et al., 2002; Dellavalle et al., 2011). This strain is characterized by the inactivation, upon tamoxifen injection, of TGF-β Receptor 2 (TGF-βR2) in TNAP expressing cells. We first verified by immunofluorescence whether TNAP^+^ cells express TGF-βR2 and found that virtually all TNAP^+^ cells are also positive for TGF-βR2, both in Sham and Ang II treated mice (Fig. 3a, b). We then injected the *Tnap*^*cre*^;*Tgfβr2*^fl/fl^ transgenic mice with 100mg/kg of tamoxifen (TAM) once a day for 5 days; at day 7 we implanted subcutaneous mini-pumps releasing 2.8mg/kg per day of Ang II for 2 weeks (Fig. 3c). We verified Cre recombination efficiency on the total heart of the *Tnap*^*cre*^;*Tgfβr2*^fl/fl^ mice by RT-qPCR for the exon 4 of the *Tgf-βr2*, the exon excised by Cre recombinase (Chytil et al., 2002), and found approximately 50% reduction in its expression in *Tnap*^*cre*^;*Tgfβr2*^fl/fl^ compared to control (Fig. 3d). We confirmed this recombination efficiency by counting the number of TNAP/tdTomato double positive cells over the total number of TNAP^+^ cells in the *Tnap*^*cre*^;*Rosa26-tdTomato* mice. Injection of control oil, did not modify the expression of TNAP.

**Figure 3.**
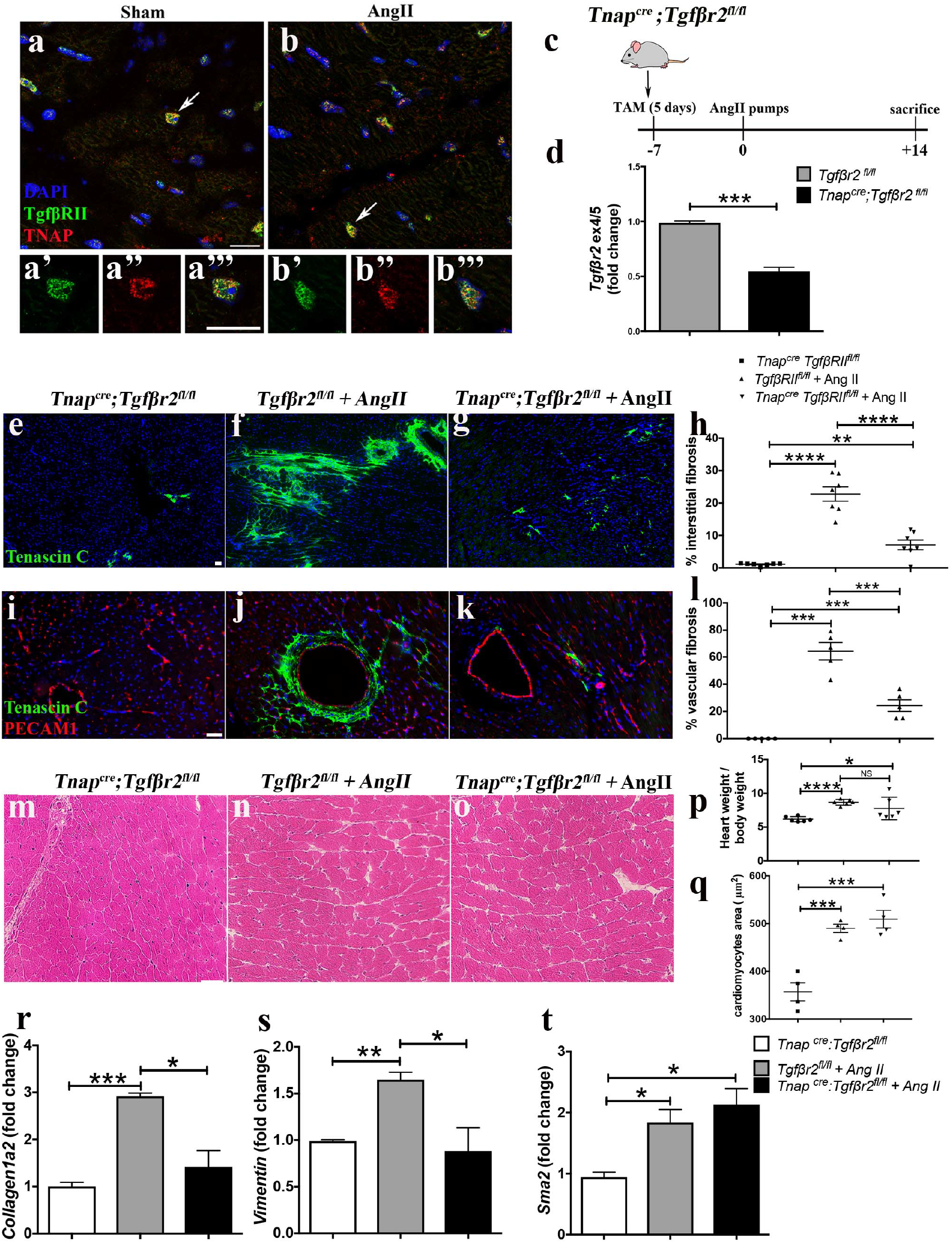
Deletion of Tgf-βr2 in TNAP positive cells reduces Ang II induced fibrosis in the heart. (a, b) Confocal images of sections from Sham and Ang II treated heart labelled for TNAP and TGF-βRII. Arrows indicates double positive cell in the ventricle. High magnification images for separate channels shown below (a’-b’’’). (c) Schematic diagram of the experimental protocol. (d) RT-qPCR on Tgf-βr2^fl/fl^ and Tnap^cre^;Tgf-βr2^fl/fl^ heart ventricle extracts for the detection of Tgf-βr2 exon 4/5 (mean ± S.D., n=3). Immunofluorescence for Tenascin C (e-g) and Tenascin C and PECAM1 (i-k) on Tnap^cre^;Tgf-βr2^fl/fl^ without Ang II, Tgf-βr2^fl/fl^ with Ang II and Tnap^cre^;Tgf-βr2^fl/fl^ with Ang II heart ventricles sections. Percentage of interstitial (h) and vascular (l) fibrosis for the three groups. The quantification has been done on 6 different sections for each sample (mean ± S.D., n=7 and n=5 for each group, respectively). Hematoxylin/eosin staining on heart sections from Tnap^cre^;Tgf-βr2^fl/fl^ without Ang II (m), Tgf-βr2^fl/fl^ with Ang II (n) and Tnap^cre^;Tgf-βr2^fl/fl^ with Ang II (o). (p) Heart and body weight ratio for the three groups. (q) Quantification of cardiomyocyte diameter for the three groups (mean ± S.D., n=4). (r-t) RT-qPCR for Collagen1a2, Vimentin and Sma2 performed on Tnap^cre^;Tgf-βr2^fl/fl^ without Ang II, Tgf-βr2^fl/fl^ with Ang II and Tnap^cre^;Tgf-βr2^fl/fl^ with Ang II cDNA samples. Mean normalized fold changes ± S.D., n=3. Scale bars: 20 µm. Two-tailed t test was used to assess statistical significance: * p<0.05, ** p<0.01, *** p<0.001, **** p<0.0001.

We next quantified the area of fibrosis in *Tnap*^*cre*^;*Tgfβr2*^fl/fl^ Sham treated mice, *Tgfβr2*^fl/fl^ treated with Ang II and *Tnap*^*cre*^;*Tgfβr2*^fl/fl^ treated with Ang II. We found a strong reduction of interstitial and vascular fibrosis in the *Tnap*^*cre*^;*Tgfβr2*^fl/fl^ Ang II treated mice in comparison with the *Tgfβr2^fl/fl^* treated with Ang II (Fig. 3e-l). This result was confirmed by Azan-Mallory staining and by IF analysis, showing reduced expression of Collagen 1a2 and MMP1 in *Tnap*^*cre*^;*Tgfβr2*^fl/fl^ Ang II treated mice in comparison with the *Tgfβr2^fl/fl^* treated with Ang II (Supplementary Fig. 4a-i).We analysed cardiac hypertrophy in *TNAP*^cre^ *Tgf β RII*^fl/fl^ mice treated with Ang II compared with the control by calculating the heart weight/ body weight ratio and by measuring the area of cardiomyocytes, but we found no difference in cell size and in overall hypertrophy within the two groups (Fig. 3m-q), indicating that the reduction of fibrosis is not accompanied by a reduction in hypertrophy. We next characterized the expression of fibrotic genes by RT-qPCR in the three sample groups and found a significant reduction in *Collagen 1a2* and *Vimentin* expression in the *Tnap*^*cre*^;*Tgf*-*βr2*^*fl/fl*^ mice treated with Ang II compared with the controls (Fig. 3r,s), while *Sma2* expression did not change (Fig. 3t). Moreover, also immunostaining with antibodies against collagen 1 and MMP1 showed a dramatic reduction of areas of fibrosis, as also confirmed by Azan-Mallory staining (Supplementary figure 4). These data prove that TNAP is specifically expressed and upregulated in cells with active TGF-β signalling that underlie cardiac fibrosis. It is remarkable that a 50% reduction of expression in a minority of the cardiac cell population leads to a dramatic decrease in the level of fibrosis, suggesting a specific role of TNAP+ cells in regulating fibrosis.

**Figure 4.**
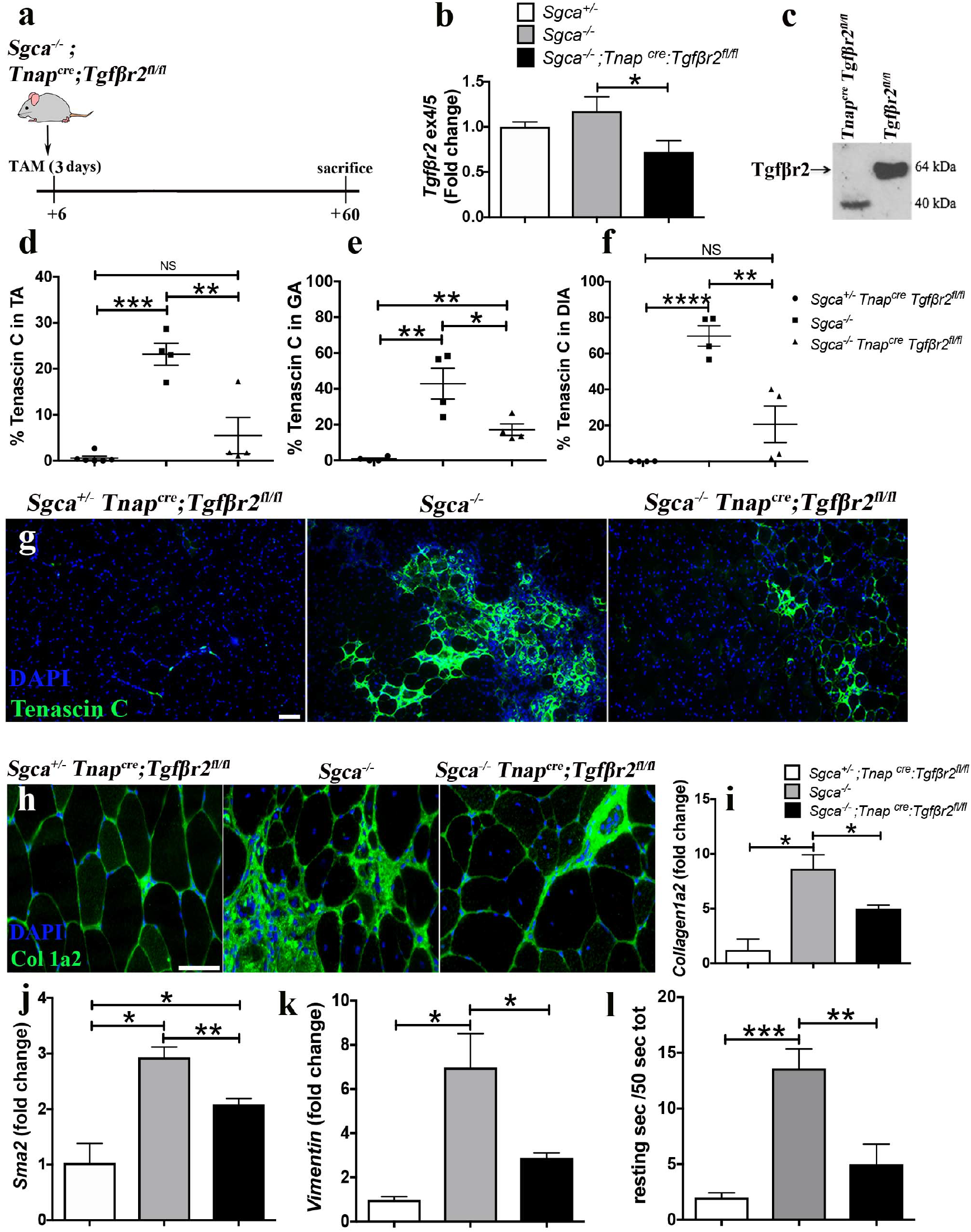
Deletion of Tgf-βr2 in TNAP positive cells reduces skeletal muscle fibrosis in Sgca^−/−^ mice. (a) Schematic diagram of the experimental protocol. (b) RT-qPCR on Sgca^+/−^, Sgca^−/−^ and Sgca^−/−^ Tnap^cre^;Tgf-βr2^fl/fl^ Tibialis Anterior for the detection of the exon 4/5 of Tgf-βr2 (mean ± S.D., n=3). (c) Western blot for Tgf-βr2 on Tgf-βr2^fl/fl^ and Tnap^cre^;Tgf-βr2^fl/fl^ proteins extracts. Quantification of Tenascin C positive areas in Sgca^+/−^, Sgca^−/−^ and Sgca^−/−^ Tnap^cre^;Tgf-βr2^fl/fl^ Tibialis Anterior (TA) (d), Gastrocnemius (GA) (e) and Diaphragm (DIA) (f) muscles. The quantification has been done on 6 different sections for each sample. Staining for Tenascin C (g) and Collagen1a2 (h) on the TA of the three groups of mice. (i-k) RT-qPCR for Collagen1a2, Sma2 and Vimentin, performed on Sgca^+/−^, Sgca^−/−^ and Sgca^−/−^ Tnap^cre^;Tgf-βr2^fl/fl^ TA cDNA samples. Mean normalized fold change ± S.D., n=3. (l) Muscle grip strength test on Sgca^+/−^, Sgca^−/−^ and Sgca^−/−^ Tnap^cre^;Tgf-βr2^fl/fl^ mice. Axis indicates the mean of immobile time over 50 second interval on upside-down wireframe (± S.D.; n=4). Scale bar represents 50 µm. A two-tailed t-test was used to assess statistical significance: * p<0.05, ** p<0.01, *** p<0.001.

Since it has been previously shown that TNAP is up regulated in dystrophic muscles (Diaz-Manera et al., 2012) we asked whether the abrogation of TGF-β signalling in TNAP^+^ cells in dystrophic mice could reduce skeletal muscle fibrosis. We took advantage of the alpha-sarcoglycan deficient (*Sgca*-null) mice, a model for Limb-Girdle Muscular Dystrophy 2D (Duclos et al., 1998) that develop severe fibrosis after 2 months of age. We generated a triple transgenic strain by crossing the *Sgca* null mice with the *Tnap*^*cre*^;*Tgf*-*βr2*^*fl/fl*^ mouse. We injected 6-7 days old pups with tamoxifen for three days and sacrificed at 2 months old (Fig.4a). We measured the recombination efficiency of Cre recombinase by RT-qPCR and western blot (Fig.4, b,c) and found it to be 50%. We next analysed the extent of fibrosis by measuring the percentage of Tenascin C positive regions and found a significant decrease in fibrosis in the Tibialis Anterior (TA), the Gastrocnemius (GA) and the Diaphragm (DIA) of the *Sgca* null *Tnap*^*cre*^;*Tgf*-*βr2*^*fl/fl*^ mice compared with the *Sgca* null controls (Fig.4 d-g). We analysed the level of Collagen1a2 by immunofluorescence (Fig.4h) and the expression level of *Collagen1a2*, *Sma2* and *Vimentin* by RT-qPCR (Fig.4 i-k). We observed a significant reduction of their expression in the *Sgca* null *Tnap*^*cre*^;*Tgf*-*βr2*^*fl/fl*^ mice compared with the *Sgca* null controls. In order to reveal whether the reduction of fibrosis in the triple transgenic mice is sufficient to improve muscle strength and function, we measured muscle grip strength by placing mice on an upside down grid (Bonetto et al., 2015). Wild-type mice started to explore the environment and spent little time in the same location. In contrast *Sgca* knockout mice spent 26% of the total time without moving, which importantly decreased to only 10% in the triple transgenic mice (Fig.4l) (n=4 each group). These data suggest that the reduction in fibrosis is associated with improved skeletal muscle function. Overall the data so far demonstrate that cells that express TNAP in fibrotic cardiac and skeletal muscle are of pivotal significance in the TGF-β-dependent development of the pathology as inhibition of TGF-β signalling in these cells is sufficient to limit fibrosis.

### TNAP inhibits SMAD2/3 phosphorylation and limits the expression of downstream fibrotic genes

Since we obtained similar results in cardiac and skeletal muscle models of fibrosis, we hypothesized that TNAP may have a role in the regulation of TGF-β signalling and that this role is conserved in both tissues. Being a phosphatase, TNAP could be involved in the dephosphorylation of p-SMAD2/3 and have thereby a regulatory role in the transcription of fibrotic genes. In order to test our hypothesis we first verified if TNAP and p-SMAD2/3 co-localize within the cell. We stained sections for TNAP and p-SMAD2/3 and saw that both of them partially co-localise in the peri-nuclear region (Fig. 5a). We confirmed their co-localization also *in vitro*. We transfected C2C12 skeletal myogenic cells with EYFP-SMAD2 and mCherry-TNAP plasmids and confirmed that the two proteins co-localize in intracellular vesicular compartments around the nucleus (Fig. 5b). Interestingly TNAP and SMAD2 co-localization was present in both Sham and Ang II treated mice (Fig. 5a). We found that TNAP not only co-localizes with SMAD2 but also physically interacts with it. We proved this by using Proximity Ligation Assay (PLA) (Shah et al., 2017) (Fig. 5c) and co-immunoprecipitation on C2C12 transfected with Myc-TNAP (Supplementary Fig. 5).

**Figure 5.**
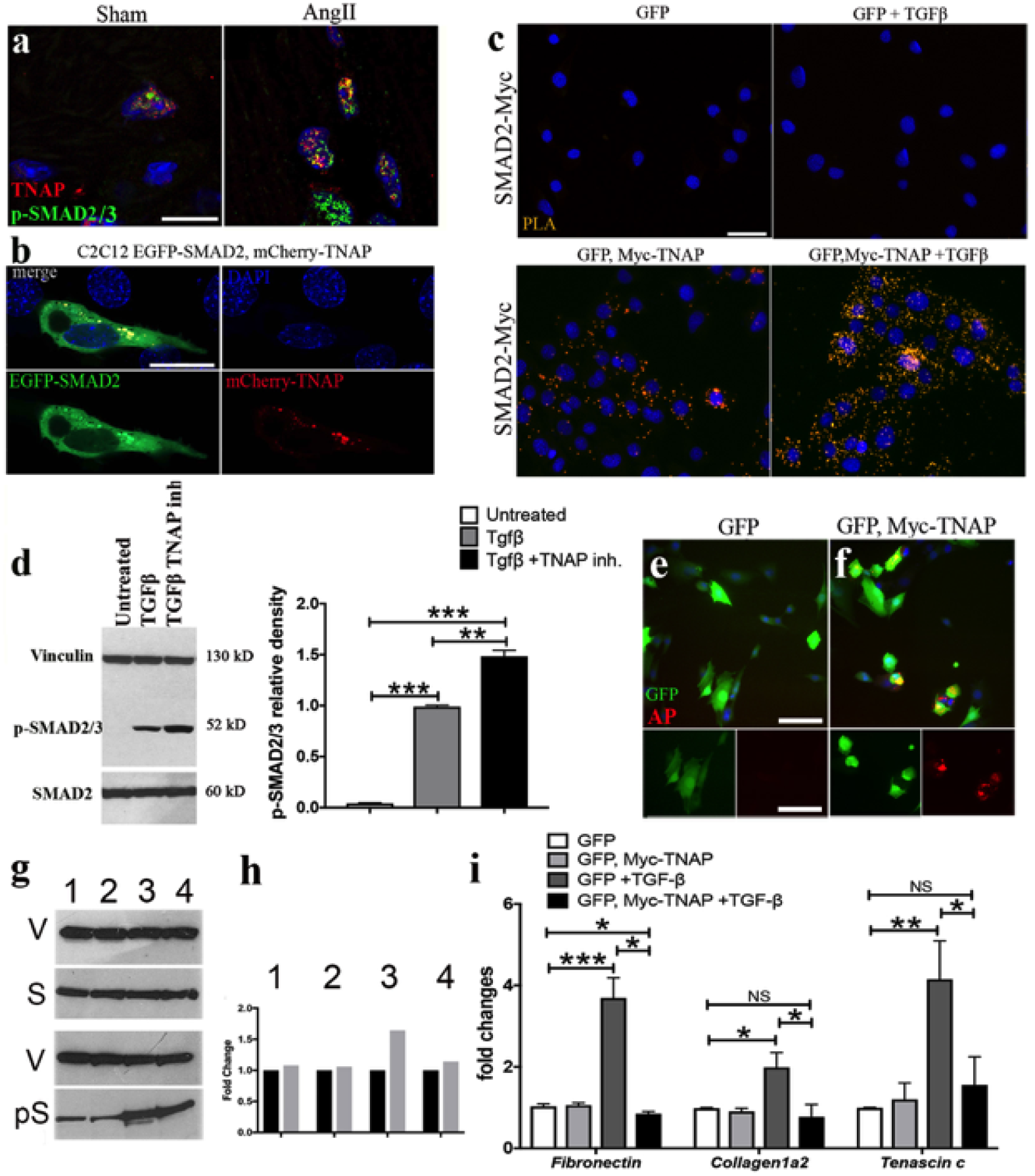
TNAP regulates SMAD2/3-dependent TGF-β signaling. (a,) Confocal images of TNAP and p-SMAD2/3 co-localization in Sham and Ang II treated heart sections. (b) Confocal images of intracellular co-localization of TNAP and SMAD2 in C2C12 cells transfected with EGFP-SMAD2 and mCherry-TNAP. (c) Proximity Ligation Assay (PLA) analysis of Myc-TNAP interaction with endogenous SMAD2 in C2C12 cells, untreated or treated with 5 ng/ml of TGF-β1 for 3h; GFP was used as transfection control (n=3 independent experiments). (d) Western blot analysis for SMAD2 and p-SMAD2/3 on cell lysate from primary cardiac fibroblasts treated with either 5 ng/ml of TGF-β1 or 5 ng/ml of TGF-β1 and TNAP inhibitor MLS-0038949 (200nM) for 1h; untreated cells were used as control. TNAP inhibition increases SMAD2/3 phosphorylation. (e, f) Immunofluorescence analysis for GFP and TNAP activity in C2C12 cells transfected with Myc-TNAP and GFP plasmids. C2C12 transfected only with GFP were used as control. (g) Western blot analysis for SMAD2 and p-SMAD2/3 on C2C12 cells transfected with Myc-TNAP and GFP plasmids, treated or untreated with 5 ng/ml TGF-β1 for 3h; C2C12 transfected only with GFP plasmid were used as control. Myc-TNAP overexpression leads to reduced SMAD phosphorylation. (h) Densitometric analysis of the Western Blot shown in (g). Grey bars: phopho-SMAD; Black bars: SMAD. Each bar represent the value normalised to Vinculin and plotted in relation to control SMAD, arbitrarily set at 1. (i) RT-qPCR for Fibronectin, Collagen1a2 and Tenascin C performed on cDNA samples from C2C12 transfected with Myc-TNAP and GFP plasmids treated with either 5 ng/ml of TGF-β1 for 3h or untreated; C2C12 transfected only with GFP were used as control. Normalized fold changes (mean ± S.D.) are shown in h (n=3 independent experiments). Scale bars: 20 µm. Two-tailed t-test was used to assess statistical significance: * p<0.05, ** p<0.01, *** p<0.001.

Our results show that endogenous SMAD2 interacts with Myc-TNAP. Notably, the TNAP-SMAD2 interaction not only occurs upon TGF-β stimulation, but also, at lower level, under control conditions. In order to understand if this interaction has a functional relevance on TGF-β signalling we treated cardiac fibroblast with TGF-β1 (5ng/ml) and the specific TNAP inhibitor MLS-0038949 (Crowley et al., 2006) (200nM) for 1 hour and analysed SMAD2/3 phosphorylation by western blot. The cells treated with TGF-β1 and TNAP inhibitor showed a significant increase in SMAD2/3 phosphorylation compared with cells treated with only TGF-β1 (Fig.5d) (n=3 independent experiments), suggesting that the blockade of TNAP leads to enhanced TGF-β signalling. To provide further evidence for TNAP regulation of SMAD2/3 activity we overexpressed plasmid encoding for Myc-TNAP in C2C12 skeletal muscle cells and stimulated the cells with TGF-β1 (5ng/ml). We chose this cell line because it has no detectable TNAP expression (Fig. 5e). We verified that the plasmid was functional by staining C2C12 with a functional assay for AP (Fig. 5e,f). Upon TGF-β1 stimulation we detected a decrease in SMAD2/3 phosphorylation in Myc-TNAP overexpressing cells compared with the GFP control (Fig. 5g). Inhibition of SMAD2/3 phosphorylation was not dramatic because on average approximately 20-30% of C2C12 cells are transiently transfected. Together these data prove that TNAP is capable and in itself sufficient to limit SMAD2/3 phosphorylation. We next analysed the expression of fibrotic genes downstream p-SMAD2/3 by RT-qPCR and observed that *Fibronectin1*, *Collagen1a2* and *Tenascin C* expression were downregulated in Myc-TNAP overexpressing cells compared with the control (Fig. 5h), indicating that TNAP induced dephosphorylation of p-SMAD2/3 is sufficient to limit fibrotic genes transcription. Taken together these data indicate that TNAP regulates SMAD2/3 phosphorylation levels and is a direct attenuator of cellular pro-fibrotic signaling pathways.

## Discussion

Despite ever increasing understanding of the molecular processes that trigger fibrosis across different organs there are only few therapies available and most of them only indirectly target fibrotic processes (Rockey et al., 2015). TGF-β1 binding to the receptors on the plasma membrane leads to the phosphorylation of R-SMADs, their translocation into the nucleus and transcription of pro-fibrotic genes. Given the importance of TGF-β signaling and the phosphorylation of R-SMADs in fibrosis, the identification of phosphatases, able to switch off the TGF-β pathway, has been of interest in the past ten years since they could also be targets for anti-fibrotic drugs. Nevertheless, until now, only one phosphatase, PPM1A, has been identified (Lin et al., 2006). Here we show that TNAP is able to dephosphorylate SMAD2/3 and consequently limit the transcription of pro-fibrotic genes.

An abundant literature exists on TNAP function in bone development and calcification (Byon et al., 2008; Millan, 2013); yet the role of TNAP in the heart has only been partly elucidated. TNAP overexpression has recently been associated with increased cardiac fibrosis and vascular calcification. Sheen et al. demonstrated that the induced expression of TNAP in vascular smooth muscle cells is sufficient to cause medial vascular calcification (Sheen et al., 2015). Similarly Romanelli and colleagues showed that TNAP overexpression in vascular endothelium in mice leads to arterial calcification and subsequent coronary atherosclerosis (Romanelli et al., 2017). These studies described the effects of sustained TNAP overexpression, but the role of TNAP under physiological levels of expression has not been addressed. Indeed, it is counterintuitive that tissues where calcification is exclusively pathological, have maintained TNAP expression during evolution. This suggests that TNAP may also play beneficial roles in striated muscle that is eventually overridden by sustained fibrotic stimuli; however this role has thus far remained unknown. Here we analysed two different models of fibrosis: rapid-onset cardiac fibrosis induced by the hypertensive molecule Ang II, and chronic skeletal muscle fibrosis in the *Sgca* null mice. We confirm earlier evidence showing that TNAP is upregulated in tissue fibrosis (Herencia et al., 2015) (Gan et al., 2014), but also show here that the number of cells expressing TNAP increases in both heart and skeletal muscle. Furthermore, as the abrogation of TGF-β signaling in TNAP positive cells dramatically reduces vascular and interstitial fibrosis, we can conclude that TNAP is actively expressed in cells driving tissue fibrosis. Since in the heart, not only interstitial/perivascular cells but also endothelial cells and cardiomyocytes participate to the process (though likely to a minor extent), it is interesting to note that TNAP has a similar pattern of expression, being more abundant in interstitial/perivascular cells but also present in the other cell types of the heart.

TGF-β signaling relies on trafficking of activated phosphorylated R-SMAD proteins from the plasma membrane to the nucleus (Chen, 2009; Hough et al., 2012; Nakao A, 1997). SMAD2/3 inactivation during this process would enable a cell to rapidly limit the expression of pro-fibrotic genes even in the presence of TGF-β secreted in the inflammatory tissue environment. Here we show that TNAP inhibition enhances SMAD2/3 phosphorylation whereas TNAP overexpression has the opposite effect. We demonstrate that TNAP interacts with SMAD2. Our data collectively prove that TNAP is an efficient inhibitor of SMAD2/3 and limits SMAD-dependent transcription of pro-fibrotic genes. Intriguingly, TNAP interacts with SMAD2 even in cells that are not exposed to TGF-β1. These results demonstrate that TNAP likely has a housekeeping function during normal tissue growth and homeostasis, which explains its basal level expression in cardiac and skeletal muscle. In addition to the molecular mechanism described here, TNAP may also indirectly affect the activity of numerous phosphorylation-dependent pathways as this enzyme is known to regulate the level of cytoplasmic inorganic phosphate (Pi) (Hessle et al., 2002).

We show here that the ubiquitously expressed TNAP is an essential regulator of SMAD2/3-dependent TGF-β signaling in physiological conditions and in tissue fibrosis. These results provide a novel insight into the mechanisms that control the expression of extracellular matrix genes and may lead to the development of novel therapies for fibrosis.

## Material and methods

### Animals

Mice were maintained in pathogen-free conditions at Manchester University animal facility and fed standard rodent diet. Animal studies were performed according to the United Kingdom Animals (Scientific Procedures) Act 1986 and approved by the University of Manchester Ethics Committee. Angiotensin experiments were approved under Home Office license no. 40/3625 and no. P3A97F3D1. All the other experiments were approved under licence no.707435. *Sgca* null mice (Duclos et al., 1998), *Tnap*^*cre*^ mice (Dellavalle et al., 2011) and *Tgfbr2*^*flox/flox*^ (Chytil et al., 2002) mice were backcrossed onto C57BL/6 mice (Charles River Laboratories Inc, Wilmington, MA, USA) and were genotyped as previously described (Dellavalle et al., 2011). Mice were culled by administration of isoflurane and consequent cervical dislocation at appropriate time points. Hearts and muscles were collected in ice-cold PBS and then embedded in glue or OCT inclusion media. Samples were stored at −80°C before processing.

### Osmotic Minipump Infusion of Angiotensin II and Tamoxifen administration

In order to generate cardiac hypertrophy, Angiotensin II (Sigma Aldrich, St. Louis, MO, USA Cat. No. A9525) (2.8mg/kg/day) or vehicle (ddH_2_O) was administrated to 10 week old wild type (wt) and transgenic male mice via osmotic mini-pumps (1002, Alzet, Cupertino, CA, USA) implanted subcutaneously. Briefly, the mice were anaesthetized with <u>isoflurane</u> (5% induction; 2–3% maintenance) in oxygen, using a small nose cone attached to a modified Bane circuit. The skin on the dorsal surface of the animal was shaved with the clipper and sterile scissors were used to make a 0.5 cm horizontal incision. Subcutaneous tissue was spread using a haemostat to create a pocket in which the minipump was inserted. The incision was closed with absorbable suture (5-0 Maxon). After the surgery the mice received 0.1mg/kg of buprenorphine and were allowed to recover before being returned to the home cage. Hearts were isolated and analyzed for cardiac hypertrophy following 14 days of Angiotensin II infusion. In order to induce the expression of the Cre recombinase in the null *Tnap*^*cre*^ *Tgfβr2*^*fl/fl*^ and *Sgca*^−/−^ null *Tnap*^*cre*^ *Tgfβr2*^*fl/fl*^, Tamoxifen (Sigma Aldrich, Cat. No. **T5648**) was dissolved in corn oil (Sigma Aldrich, Cat. No. C8267) and injected at the following concentrations and time points: 0.3 mg/mouse in 6 days old pups once a day for 3 consecutively days and 3 mg/mouse in 10 weeks old mice once a day for 5 consecutively days.

### Grip test

The four limb-hanging test (also known as Kondziella’s inverted screen test) (Bonetto et al., 2015) represents a method to assess muscle strength using all four limbs and to determine the general condition over time. In this study, we modified this method to improve its variability and consistency. Briefly we placed the mice upside down on a grid (50cm x 30cm) and we counted the number of seconds in which the mice were not moving on the grid, over a total period of 50 sec each mice. We calculated the resting time on the grid as the average of 3 different tests for each mouse, and we analysed 4 mice per each group.

### PLA and Immunofluorescence

PLA was performed according to the manufacturer’s instructions using the Duolink In Situ Orange Kit Goat/Rabbit (Sigma Aldrich, Cat. No. DUO92106). Immunofluorescence was performed as previously described (Arno et al., 2014). Briefly sections (5 µm) were fixed in 4% Paraformaldehyde (Sigma Aldrich, Cat. No. P6148) in PBS, pH 7.2 for 20 min; then washed 3 times, 5 min each, in PBS and the blockage of non-specific binding was done by using the following blocking buffer: PBS 1X/Fetal bovine serum (FBS) 10%/Bovine serum albumin (BSA) 1 mg/ml/Triton X 100 0.1%, for 1 h at room temperature. In all steps slides were kept in humidified chambers. For the TNAP staining the sections were additionally fixed with methanol-acetone for 10 min. Antibodies were diluted in blocking buffer and incubated at +4°C overnight as suggested by manufacturer’s instructions. The following day, sections were rinsed in PBS for 5 min, 3 times, and fluorescent secondary antibodies, (Alexafluor conjugated, Thermo Fisher Scientific) diluted in blocking buffer were applied accordingly manufacturer’s instructions. Slides were washed 3 times in PBS for 5 min and incubated in DAPI (Sigma Aldrich, Cat. No.10236276001) for nuclei counterstaining. The following antibodies were used: goat α-TNAP (Santa Cruz Biotechnology, Dallas, Texas, USA Cat. No. c-23430) 1:100; rabbit α-TNAP (Abgent, Suzhou city, China. Cat. No. AP1474C) 1:100; rat α-PECAM (Dev. Studies Hybridoma Bank) 1:3; rabbit α-Laminin (Santa Cruz Biotechnology Cat. No. sc-17810) 1:100: chicken α-GFP (Abcam, Cambridge, UK Cat. No. ab13970) 1:500; rabbit α-collagen1a2 (Merck Millipore, Billerica, MA, USA) 1:75; rabbit α-Tenascin C (Merck Millipore Cat. No. AB19013) 1:150; chicken α-Vimentin (Novus biologicals, Littleton, Colorado, USA Cat. No. NB300-223) 1:300; rabbit α-TGF-βRII (Novus biologicals Cat. No. NB100-91994) 1:100; rabbit α-SMAD2 (Cell Signalling, <u>Danvers, Massachusetts</u>, USA Cat. No. D43B4) 1:100; rabbit α-pSMAD2/3 (Cell Signalling Cat. No. D27F4) 1:100; rabbit α-Ki67 (Abcam Cat. No. ab15580) 1:200. Appropriate fluorophore-conjugated secondary antibodies (Alexa-Fluor 488 nm, 594 nm and 647 nm, Molecular Probes) were applied accordingly manufacturer’ instruction. Nuclei were visualized using DAPI. For the detection of TNAP activity, the Red Alkaline Phosphatase (Red AP) Substrate Kit (VECTOR Laboratories, Peterborough, UK SK-5100) was used according to manufacture instructions. Light (Zeiss Axio Imager M2, with x5, x10 and x20 objectives) and confocal (Leica, SP5 with x40 and x63 objectives) microscopy was performed to analyze tissue and cell staining. For confocal imaging, pictures were acquired at 1× line average and 3× frame average on a single confocal plane. Analysis was performed by using Zeiss ZEN 2 software and Adobe Photoshop CS5 Version 12.0×64 software. Quantification of Tenascin C^+^ regions in hearts and skeletal muscles were done by using ImageJ 2.0.0-rc-9.

### Cell culture

C2C12 cells from European Collection of Authenticated Cell Cultures, Public Health England (Porton Down, UK) were cultured in DMEM (Gibco, ThermoFisher, Waltham, MA, USA), supplemented with 1% L-Glutamine, 10% fetal bovine serum (FBS), 1% penicillin-streptomycin (ThermoFisher Cat. No. 15140122). Cells were transfected with Lipofectamine 2000 (Thermo Fisher Scientific Cat. No. 11668027) according to manufacture instructions. The following plasmids were used for transfection: pEGFP-N1 (Clontech Cat. No. 6085-1), pCDNA3.1 N-Myc-TNAP and pCDNA3-mCherry TNAP. In details Smad2/TNAP ORF was RT-PCR amplified from mouse cDNA. PCR products were inserted into EcoRI/XhoI of the EYFP/mCherry/myc-pcDNA3 vectors. Cardiac fibroblasts were obtained from neonatal and adult C57Bl/6 mice. Mice were killed by cervical dislocation and their hearts were rapidly removed. The hearts were then mashed with scalpels and digested with 10□ml collagenase solution (120□mg collagenase A (Roche Cat. No. 10 103 578 001) and 12□mg protease (Sigma Cat. No. 10165921001) dissolved in 80□ml PBS solution) at 37□°C for 15□min, for five times. Cells were collected after each digestion and the collagenase was deactivated by addition of 50% FBS. The fibroblasts were centrifuged for 5□min at 220□g. The cell pellet was resuspended in 10□ml ACF media (80% DMEM, 20% FBS, 1% penicillin/streptomycin and 1% non-essential amino acids). Fibroblasts were then plated in 10□ml delta-NUNC tissue culture plates (ThermoFisher Cat. No. 150318) overnight. The next day, the media was removed and replaced with 10□ml ACF media. C2C12 and cardiac fibroblasts were serum starved for 24 h before adding medium containing 5 ng/ml recombinant TGFβ1 for 2 days (R&D Systems, Minneapolis, USA Cst. No. Cat. No. 7666-MB-005).

### Western Blot and Immunoprecipitation

Hearts and skeletal muscles from wt, TNAP^cre^ TGF-β RII^fl/fl^ and alpha-sarcoglycan null TNAP^cre^ TGF-β RII^fl/fl^ and controls were homogenized with the homogenizer SHM1 (Stuart Equipment Stone, Staffordshire, UK Cat. No. 1171631) in RIPA lysis buffer (50 mM Tris HCl, pH 8.0, 150 mM NaCl, 1% NP-40, 0.5% sodium deoxycholate, 0.1% SDS, supplemented with Phosphatase and Protease inhibitor cocktail, Sigma Aldrich Cat. No. PCC 1010). Cold lysis buffer (RIPA) was added into the wells after removing the media and washing the cells with PBS. The cells were scraped into sterile, pre-cooled tubes. The samples were incubated (with rotation) for 30-60 min at 4°C. Soluble proteins were quantified with Bradford Protein Assay Kit (Bio-Rad Cat. No. 5000006). 50 µg of protein extract was run in SDS-polyacrylamide gel electrophoresis (PAGE) and subsequently blotted on a nitrocellulose membrane. For immunoprecipitation experiments C2C12 cells were collected in lysis buffer (50mM Tris-HCl pH 7.8, 150mM NaCl, 1mM CaCl_2_ and 1% Triton X-100) supplemented with Protease inhibitor cocktail (Sigma-Aldrich Cat. No.P8340). 1 mg/ml of pre-cleared lysates were incubated with anti-Smad2 (Cell Signalling, Cat. No. D43B4) at dilution 1:50 or the negative control rabbit IgG in lysis buffer with rotation overnight at +4 °C. The samples were then incubated with protein A magnetic beads (Merc Millipore Cat. No. LSKMAGA10) for 1h at +4 °C (with rotation) and the immuno-complexes were detected by western blot. The following primary antibodies were used: goat α-Myc (1:1000, Abcam Cat.No. ab9132); rabbit α-TGF-βRII (Novus biologicals Cat. No. NB100-91994) 1:500, mouse α-Vinculin (Santa Cruz Biotechnology Cat. No. H300) 1:6000, rabbit α-SMAD2 (Cell Signalling Cat. No. D43B4) 1:1000; rabbit α-pSMAD2/3 (Cell Signalling Cat. No. D27F4) 1:1000. Secondary antibodies (Dako Donkey anti-goat P0449; Dako Goat anti-rabbit P0448) were incubated for 1 h at RT, and signals were revealed using Millipore ECL kit (Cat. No. WBKLS0500). For quantitative measurement, blots were analyzed with ImageJ 2.0.0-rc-9, normalizing band intensities to Vinculin levels.

### Real time and standard RT-PCR

Total RNA from heart, skeletal muscle and cells was extracted by homogenizing the samples in Trizol (Thermo Fisher Scientific, Cat. No. 15596018) reagent following the standard manufacturer’s procedure. Chloroform was added to the samples that were incubated for 15 min at room temperature and then centrifuged at 12000g for 15 min. The aqueous layer was mixed with isopropanol, incubated for 10 min and centrifuged for 10 min at 12000g. 75% of ethanol was added and the samples were centrifuged at 12000g for 15 min. After the elimination of ethanol the RNA was dissolved in 50 ul of H20. DNase (Thermo Scientific, Cat. No. 18068015) digestion was done on all the samples. RNA was quantified with Nanodrop 2000 and equalized to same concentration before cDNA synthesis. cDNA synthesis was performed using ThermoScript RT-PCR System (Thermo Scientific, Cat. No. K1622) according to the manufacturer’s instructions in final volume of 20 µl. RT-qPCR was carried out using FastStart Essential DNA Master Mix (Cat. No. 06402712001; Roche diagnostics GmbH, Mannheim, Germany). RT-qPCR was performed in five biological replicate, using standard dilution series on every plate. Samples were normalized by using the housekeeping genes *Gapdh* and *Rlp19* with the following primers:

Gapdh F: 5’-AGGTCGGTGTGAACGGATT-3’, Gapdh R: 5’-TGTAGACCATGTAGTTGAG-3’ and Rpl19 F: 5’-ATGAGTATGCTCAGGCTACAGA-3’, Rpl19 R: 5’-GCATTGGCGATTTCATTGGTC-3’.

Specific primers were used for gene expression analysis:

*Tenascin C*: F5’-ACGGCTACCACAGAAGCTG 3’; R5’-ATGGCTGTTGTTGCTATGGCA 3’

*Fibronectin1*: F5’-TTCAAGTGTGATCCCCATGAAG 3’; R5’-CAGGTCTACGGCAGTTGTCA 3’

*Vimentin*: F5’-CGGCTGCGAGAGAAATTGC 3’; R5’-CCACTTTCCGTTCAAGGTCAAG 3’

*TGF-βr2* exon4/5: GACCTCAAGAGCTCTAACATCC3’; R5’-CTAGAACTTCCGGGGCCATG3’

*TGF-βr2:* F5’-CCGCTGCATATCGTCCTGTG3’; R5’-AGTGGATGGATGGTCCTATTACA3’

*Sma2*: F5’-5’CCCAGACATCAGGGAGTAATGG3’; R5’TCTATCGGATACTTCAGCGTCA3’

*Col1a2*: F5’GGAGAGAGGAGTCGTTGGAC-3’; R5’GTTCACCCTTCACACCCTGT3’

### Statistics

Data are expressed as the mean ± standard deviation (SD) or standard error of the mean (S.E.M.) of independent experiments. Comparisons were made using the unpaired t-test. Statistical tests were carried out using PRISM6.0e (GraphPad Software, La Jolla, CA). A p-value less than 0.05 was considered statistically significant.

